# eDAVE - extension of GDC Data Analysis, Visualization, and Exploration Tools

**DOI:** 10.1101/2023.09.13.557577

**Authors:** Jan Bińkowski, Olga Taryma-Leśniak, Katarzyna Ewa Sokolowska, Patrycja Kamila Przybylowicz, Melanie Staszewski, Tomasz Kazimierz Wojdacz

## Abstract

Publicly available repositories such as Genomic Data Commons or Gene Expression Omnibus are a valuable research resource useful for hypothesis driven research as well as validation of the results of new experiments. Unfortunately, frequently advanced computational skills are required to mine those repositories making the use of those opulent research resources challenging, especially for researchers without bioinformatics expertise. To address this challenge, we have developed eDAVE, a user-friendly, web and desktop interface enabling intuitive and robust analysis of almost 12 000 methylomes and transcriptomes from over 200 types of cells and tissues deposited in the Genomic Data Commons repository.

Web app implemented in Python, is supported for major browsers and available at: https://edave.pum.edu.pl/

## 1. Introduction

There is no doubt that epigenetic changes, such as aberrations of DNA methylation, and alterations of gene expression induced by those changes, play a key role in the pathology of disease (1). Moreover, the significance of disease-related methylation changes as biomarkers applicable in personalized medicine is rapidly increasing, as best exemplified by the fact that the newest liquid biopsy early cancer detection tests target methylation biomarkers (2). With growing interest in a translational aspect of disease specific methylation changes, large amounts of methylomics and transcriptomics data are generated and deposited in publicly available repositories, such as Gene Expression Omnibus (GEO) or Genomic Data Commons (GDC). These repositories are a valuable research resource, but deposited data are frequently unstructured and poorly integrated, making mining of these datasets challenging without appropriate bioinformatics expertise and IT infrastructure necessary for big data processing. Tools such as GEO2R (3), GEOexplorer (4), TCGAbiolinks (5) or GDC Data Analysis, Visualization, and Exploration Tools (DAVE) (6) have been developed to facilitate integration of data from these repositories but the use of these tools still requires a certain level of bioinformatics skills or is limited to only certain parts of deposited data.

To address the general limited access to bioinformatics expertise that hampers exploration of big data repositories we have developed eDAVE, a platform that enables fast and intuitive interrogation of transcriptomics and methylomics data deposited in the GDC.

## 2. Materials and Methods

### 2.1 Data curation

The data transfer between eDAVE and the GDC database utilizes two distinct communication channels: automatic programming interface (API) and GDC downloading tool. The samples list along with the clinical description is extracted with API, however the API-based communication is not sufficiently efficient to transfer large files, therefore, to transfer methylation or expression profiling data, eDAVE uses the GDC data downloading tool. To avoid the communication issues and reduce the time of processing during real-time data analysis, eDAVE is based on a local, regularly updated data repository and the repository is comprising of methylomics and transcriptomics datasets processed as described in GDC documentation (https://docs.gdc.cancer.gov/Data/Introduction/). The sample groups in the repository are annotated to distinct categories based on clinical information available in GDC, including sample type, primary diagnosis, and tissue or organ of origin. For example, category named “Primary Tumor_Kidney_Papillary adenocarcinoma” refers to profiles generated for primary tumor samples of kidney papillary adenocarcinoma. The web version of eDAVE implements some limitations due to maximum and minimum size of single sample category. Firstly, eDAVE constrains maximum number of samples (n=50, randomly selected) per single sample category due to the limited computational power of the local facilities. Secondly, eDAVE assume that categories with less than 5 samples are too small to be representative therefore are not included in the local data repository. The desktop version of eDAVE does not have those limitations and minimum/maximum thresholds can be modified by user (precise instruction is described in documentation deposited in the GitHub repository, see section 2.8).

### 2.2 eDAVE modules

The data analysis in eDAVE is based on four modules: differential features explorer, single probe/gene explorer, methylation-expression association explorer and cluster explorer.

### 2.3 Differential features (DEGs/DMPs) explorer

The module was designed to automate the identification of differentially expressed genes (DEGs) or differentially methylated positions (DMPs) between two categories of samples. In that module, a user choses the data type for the analysis (methylation or expression). Then the categories of samples for comparative analysis, statistical significance level (alpha), and minimum effect size for methylation analysis: |delta| (defined as absolute value of difference between average methylation levels between compared groups) or expression analysis: |log2(FC)| (defined as absolute value of log2 transformed ratio of average expression levels between compared groups). After submission of the request, the application returns a volcano plot describing the effect size metric (change of methylation or expression) versus log2-transformed FDR-corrected p-value for each compared feature (an example is shown in Figure 1), as well as a data frame with the results of the analysis for each analyzed feature including: mean methylation/expression value, fold change, log2-transformed fold change, delta, absolute delta, delta adjusted for variance (Hedges’ *g*) type of statistical test, p-value, log10-transformed p-value, FDR-corrected p-value, log10-transformed FDR-corrected p-value as well as information if the feature is DMP/DEG according to the significance level and minimum effect size chosen by the user (exemplary output in Supplementary Table 1).

**Figure 1:**
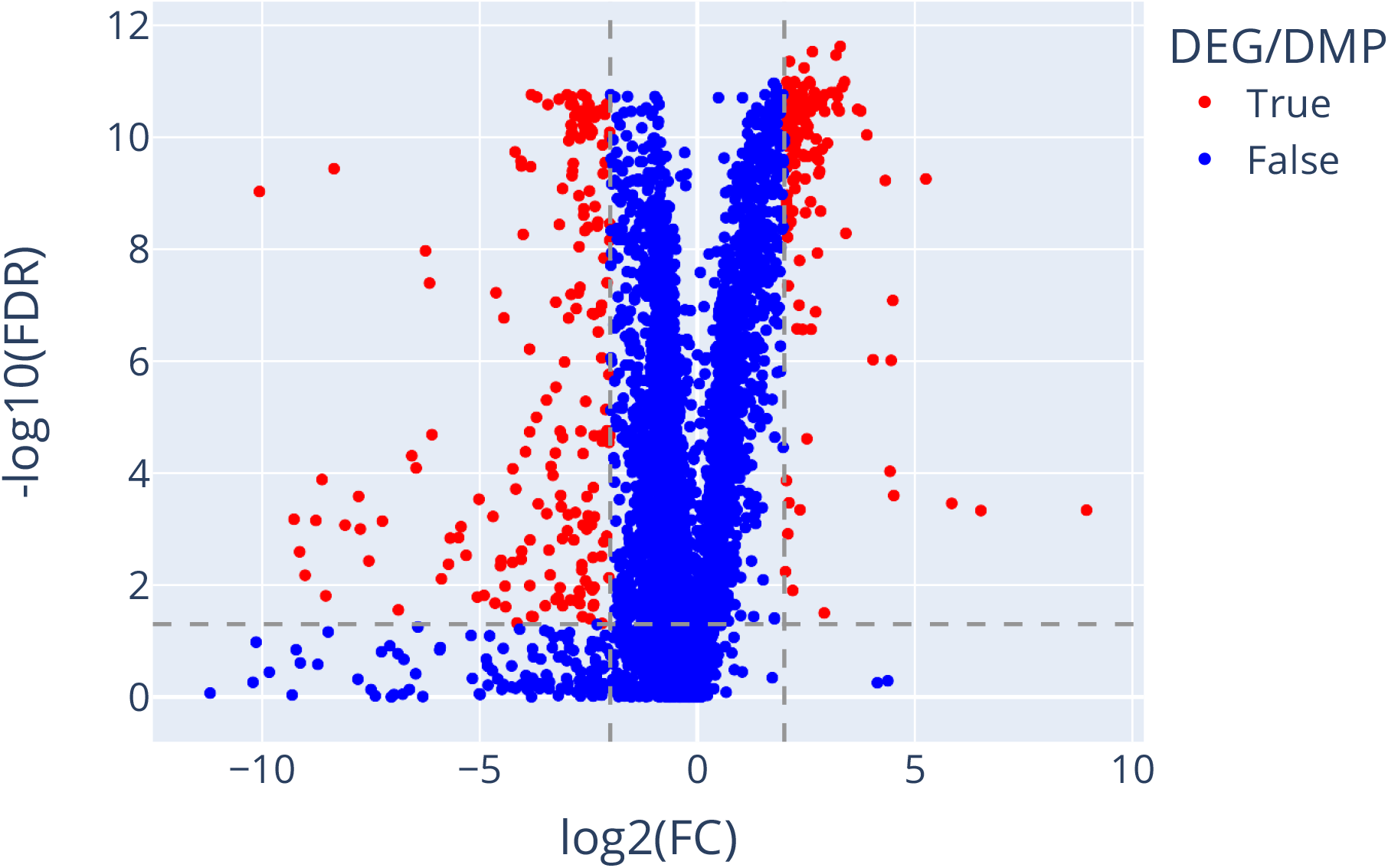
Volcano plot illustrating results of analysis of expression changes between normal breast tissue (n=50) and breast cancer infiltrating duct carcinoma (n=50). Dashed horizontal and vertical lines indicate fold-change and FDR significance thresholds respectively. Dots above these thresholds (red) indicate genes expressed differentially, while dots bellow thresholds (blue) indicate genes not meeting the significance level of differential expression between compared tissues.

### 2.4 Single probe/gene explorer

This module allows the user to compare the methylation levels at a specific CpG site or expression of a gene across multiple categories of samples. The module requires user to choose the data type (methylation or expression) for the analysis, the gene name or CpG identifier (from the Illumina EPIC or 450K manifests), optional selection of scaling method (log2, ln, log10, zero-mean and unit-variance or none), and choice of the type of visualization plot form (box, violin or scatter plots), as well as choice of statistical significance level of the analysis. After request submission, visualization of the selected variable is displayed according to the chosen plot type (an example is shown in Figure 2), along with the statistical analysis of feature differences between chosen categories of samples, including statistical parameters previously listed in section 2.3 (exemplary output in Supplementary Table 2).

**Figure 2:**
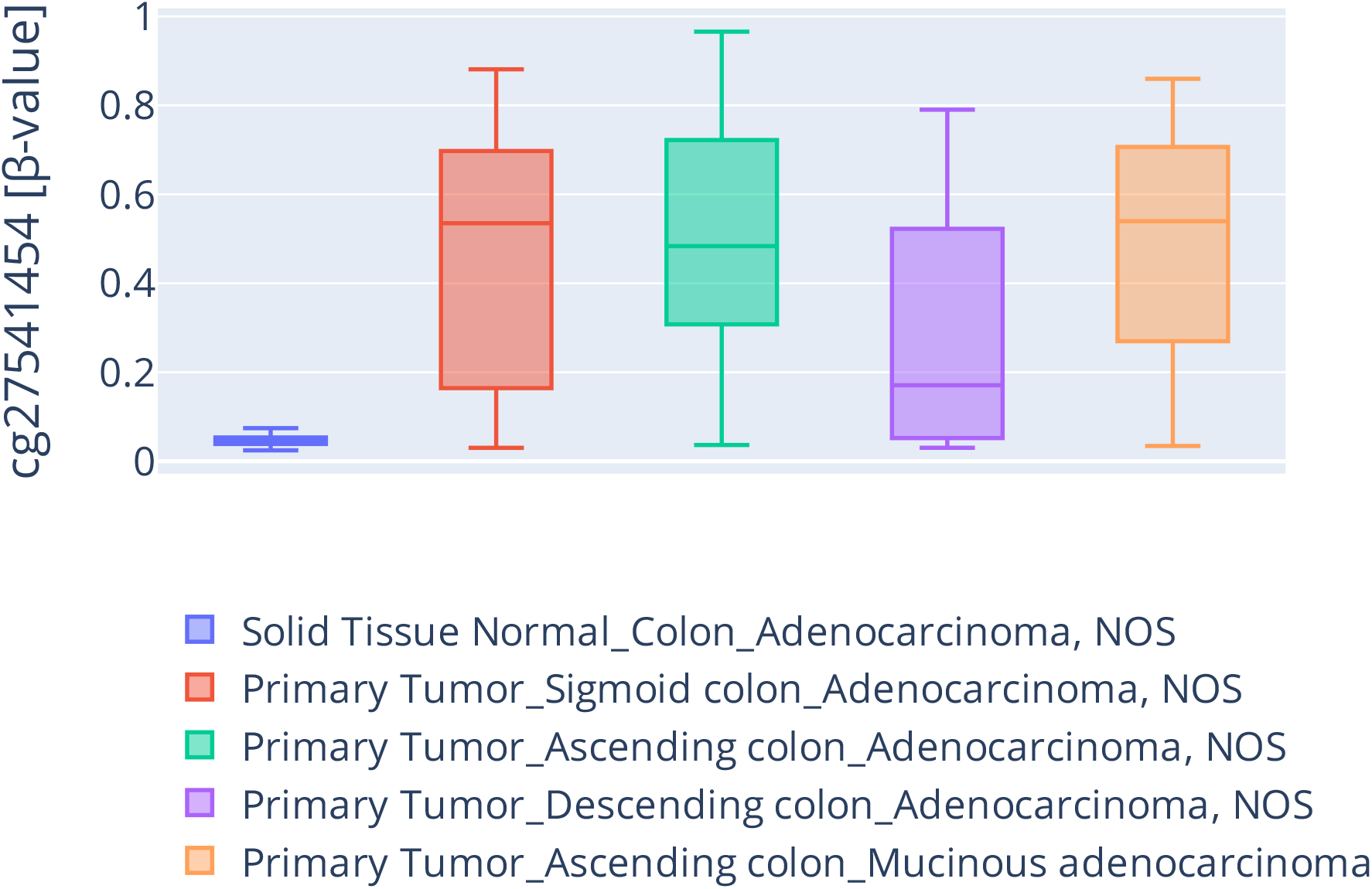
Box plot illustrating methylation levels of cg27541454 in healthy colon tissue (n=12) and four types of colon cancer: sigmoid colon adenocarcinoma (n=50), ascending colon adenocarcinoma (n=48), descending colon adenocarcinoma (n=10), and ascending colon mucinous adenocarcinoma (n=12).

### 2.5 Methylation-expression association explorer

This module computes two types of analysis describing association between the methylation levels of a specific CpG site and the expression of specified gene in a chosen sample category. The first analysis is regression-based, for which the degree of polynomial transformation can be chosen from default linear regression (degree of 1) to non-linear regression (degree > 1). The second type of analysis is bin-based, where methylation levels are converted into n-number of equally sized bins (from two to four as chosen by the user), then the expression levels of specified gene are compared between these bins. Both analyses require choice of the statistical significance for the comparisons and optional scaling method. The output of the analysis includes: regression statistics (exemplary output in Supplementary Table 3), fitted model parameters table (exemplary output in Supplementary Table 4), and plot with the regression fit (exemplary output in Figure 3). In the case of bin-based analysis, the application returns a box plot describing compared gene expression between bins (exemplary output in Figure 4), the count of samples in each bin, as well as a summary statistics table (exemplary output in Supplementary Table 5) with set of statistical parameters previously listed in section 2.3.

**Figure 3:**
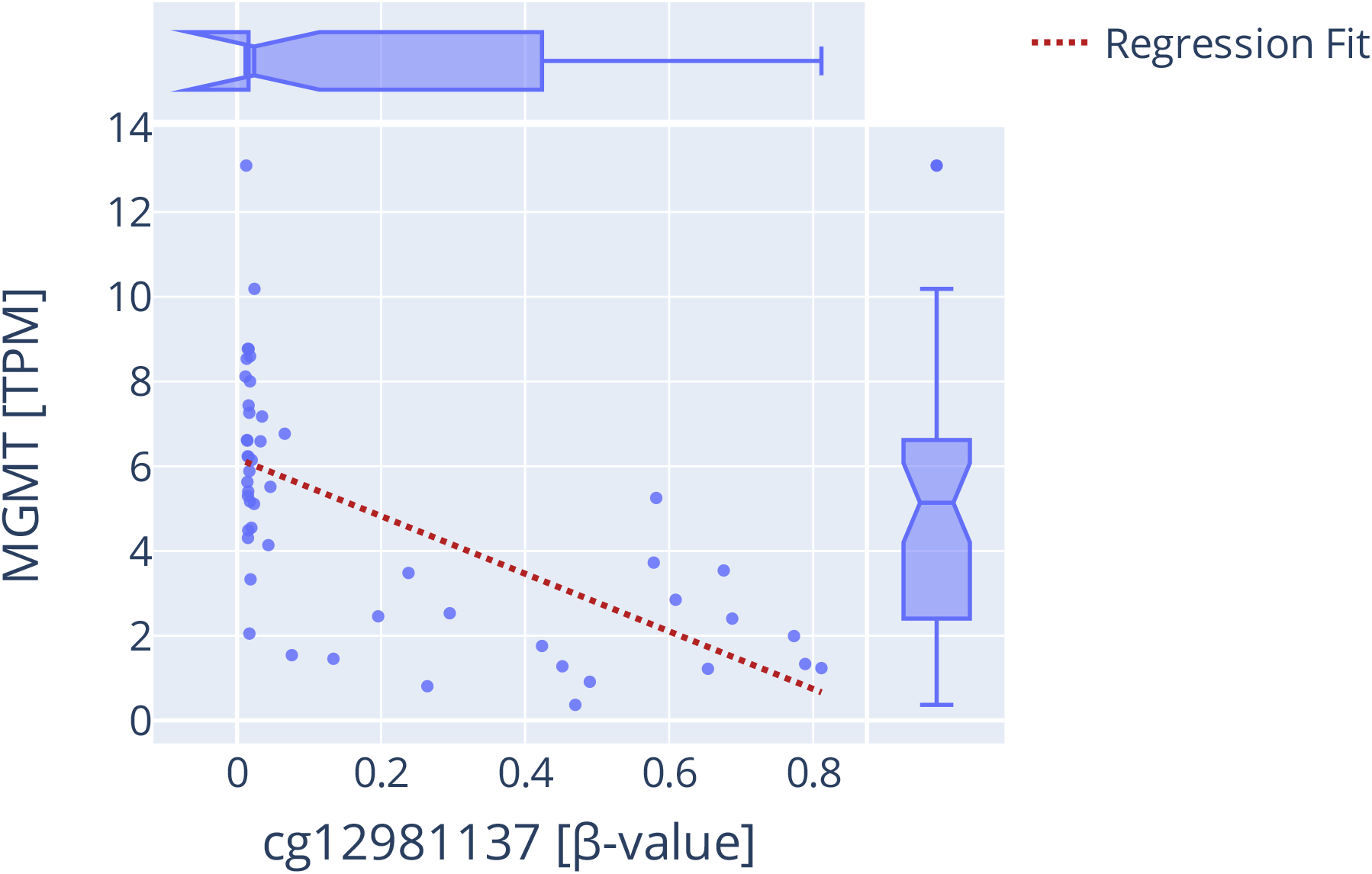
Scatter plot illustrating association between methylation of cg12981137 and *MGMT* gene expression in glioblastoma samples (n=50). Regression fit line (red dotted line) represent linear regression model fitted to data. Additionally horizontal and vertical boxplots illustrate distribution of methylation and expression levels, respectively.

**Figure 4:**
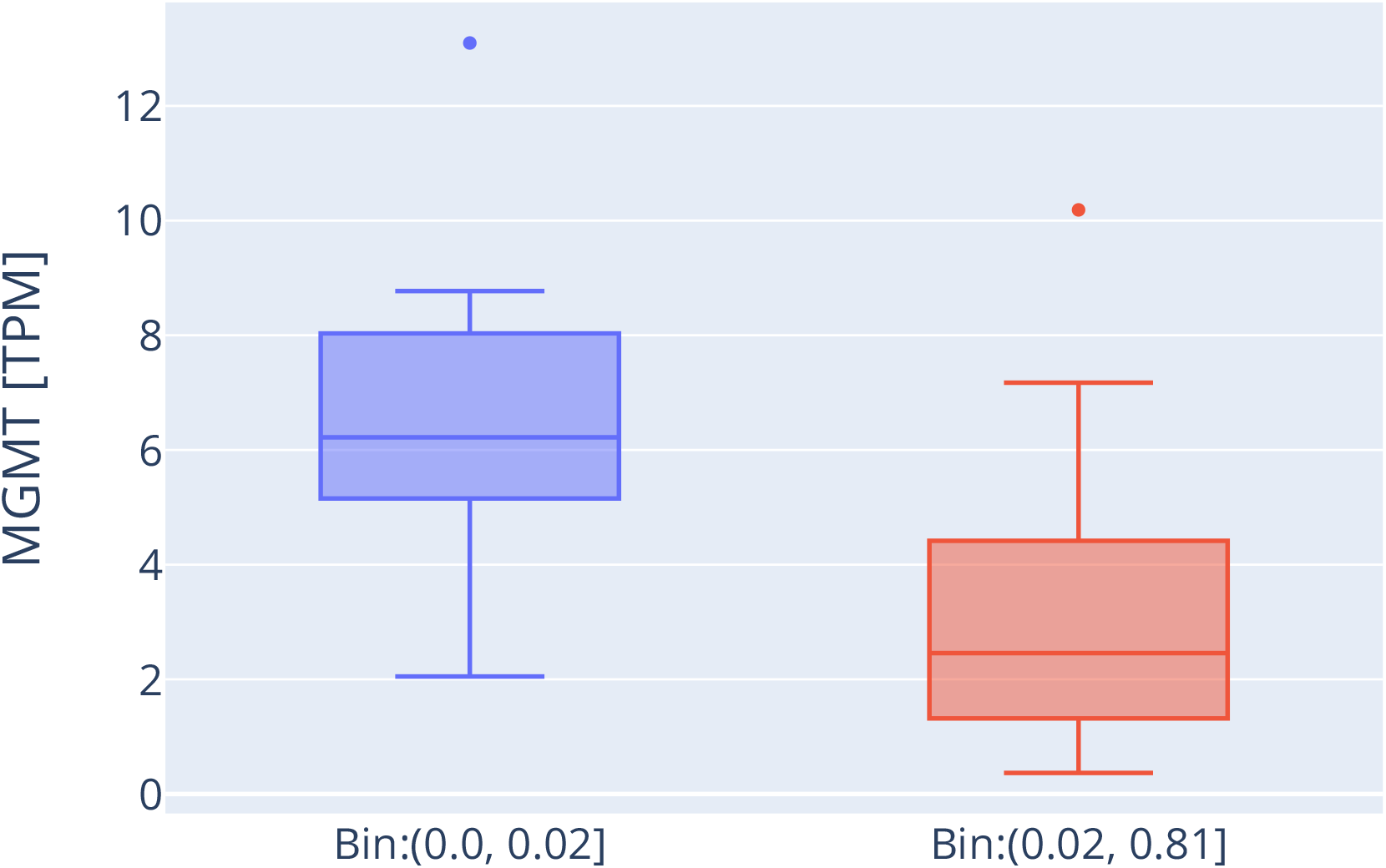
Box plot illustrating results of bin-based analysis of association between methylation of cg12981137 and *MGMT* gene expression in glioblastoma samples. First (blue) box shows expression levels in the group of hypomethylated glioblastoma samples (n=25) with methylation levels of cg12981137 in range from 0.0 to 0.02, second (red) box displays expression levels for group of hypermethylated glioblastoma samples (n=25) with methylation levels of cg12981137 in range from 0.02 to 0.81.

### 2.6 Cluster explorer

This module was designed to help identify subgroups (clusters) of similar samples within interrogated datasets of interest and to assess the discriminative power of methylation or expression changes at a subset of CpG sites or genes, respectively. Here, after selection of data type (methylation or expression), pasting the list of CpGs or genes, and selection of: samples categories, decomposition algorithm (PCA, t-SNE), and the number of dimensions to project data in (two or three). Additionally for t-SNE method perplexity parameter needs to be selected (perplexity is related to the number of nearest neighbors used in the analysis). As a result the application returns PCA or t-SNE plots. Plots illustrate the results of the cluster analysis colored by the category of sample and by the predicted cluster (exemplary output in Figure 5). The cluster prediction process is comprising five steps. Firstly, the requested dataset is scaled to mean zero and unit variance. The scaled dataset is projected into two- or three-dimensional planes using PCA or t-SNE techniques. Then, iteratively for the predefined number of clusters in a range from two to ten, the Ward clustering method based on geometric distance metric is applied. The effectiveness of the clustering for each tested number of clusters is measured using the Calinski-Harabasz (CH) metric, which by definition is higher for dense and well-separated clusters. Finally, an optimal number of clusters within the dataset is defined as a number-maximizing CH value.

**Figure 5:**
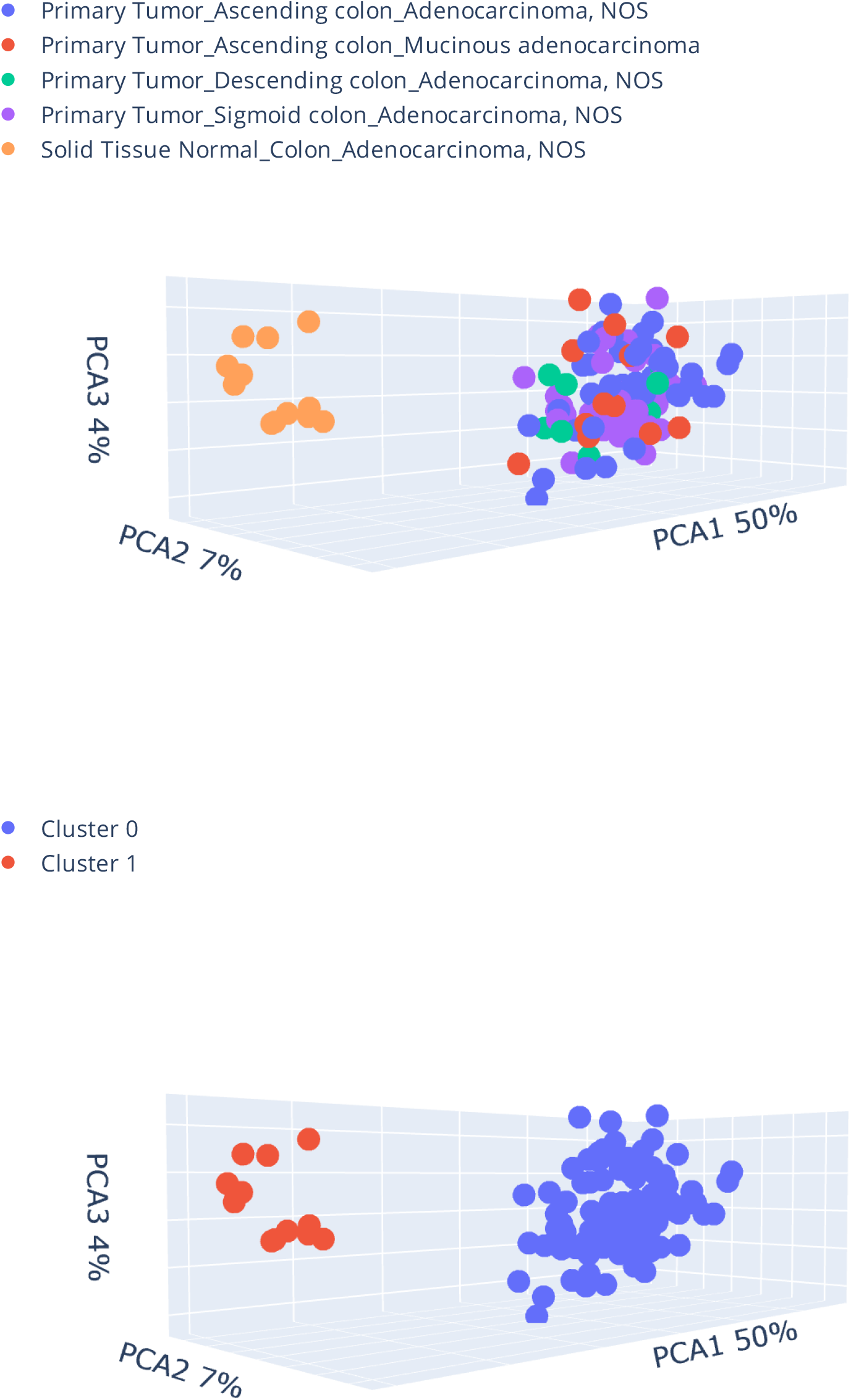
Scatter plot of 3-D PCA generated for methylation profiles of healthy colon tissue (n=12), four types of colon cancer (sigmoid colon adenocarcinoma (n=50), ascending colon adenocarcinoma (n=48), descending colon adenocarcinoma (n=10), ascending colon mucinous adenocarcinoma (n=12)) and 100 DMPs known to be discriminative for cancer colon tissue. On the upper plot, samples are marked by a category while on the bottom plot by a predicted cluster.

### 2.7 DEGs / DMPs identification procedure

The differential features (DEGs/DMPs) explorer, single probe/gene explorer, and the bin-based approach in the methylation-expression explorer share the same uniform strategy of statistical significance assessment. The procedure automatically selects the appropriate statistical test depending on the data distribution of the analyzed variable and minimizes risk of statistical assumptions violation. The procedure of statistical test selection is based on guidelines described in documentation of “pingouin” statistical library (7).

### 2.8 Results export formats

All plots generated by eDAVE include statistical information about analyzed features that can be displayed by hovering the cursor over the feature and plots can be downloaded in high resolution vector format (suitable for publication) by clicking the camera icon present in the upper-right corner of each figure. The tables with the analysis results can be exported in .csv format using “export” function. Additionally, all modules return “sample count table” with information about the number of samples in each analyzed category which can also be exported.

### 2.9 Implementation

eDAVE was implemented in Python 3.10 using Dash web framework. Python code of web app, script used to build local data repository, docker file as well as technical documentation are available from: https://github.com/ClinicalEpigeneticsLaboratory/eDAVE.

## 3. Results

### 3.1 Validation of performance of eDAVE

To verify the robustness and usability of eDAVE, we conducted a series of independent experiments and compared generated results with the results reported in previous publications.

### 3.2 Differential features (DEGs/DMPs) explorer

We first used eDAVE to perform a differential expression analysis between healthy breast tissue and breast infiltrating duct carcinoma. The platform identified a set of 369 DEGs (|log2 normalized FC| ≥ 2 and FDR ≤ 0.05, Supplementary Table 1, Figure 1), which we confirmed are altered in Breast_Mammary_Tissue (Supplementary Figure 1), and belong to categories of genes associated with breast cancer and generally soft tissue cancers (Supplementary Table 2) using FUMA platform (8).

### 3.3 Single probe/gene explorer

The hypermethylation of cg27541454 has been shown to be a potential colon cancer biomarker in both liquid and solid biopsy (9). We used eDAVE to analyze methylation levels of this CpG site in healthy colon and four different colon cancer tissues. The analysis based on the single probe/gene explorer (Figure 2, Supplementary Table 3) confirmed previously described statistically significant (FDR ≤ 0.05) elevated methylation levels in all analyzed cancerous colon tissues in comparison to healthy colon tissue.

### 3.4 Methylation-expression association explorer

It is well known that *MGMT* gene expression is regulated by methylation and the methylation of that gene is used to guide glioblastoma therapy (10, 11). We tested the association between *MGMT* promoter methylation and expression in glioblastoma samples accessible via eDAVE and confirmed a statistically significant negative association between methylation levels of cg12981137 located in the *MGMT* promoter and *MGMT* expression (Figure 3-4, Supplementary Tables 4-6).

### 3.5 Cluster explorer

We used a set of 100 CpGs (Supplementary Table 7) previously identified as being altered between colon cancer and healthy colon tissue (12) to test the accuracy of the cluster explorer. The PCA three-dimensional visualization based on those probes correctly identified two well-separated clusters of cancer and normal samples (Figure 5).

## 4. Discussion

We have developed an intuitive and robust tool for the analysis of methylation and gene expression data deposited in the GDC repository. The tool allows researchers without programming expertise easy access to the database resources comprising large amounts of methylomics and transcriptomic profiles from various types of cancer and healthy tissues. We tested the performance of eDAVE and shown its utility in both hypothesis-driven and data-validation research.

