## Supplementary figures for "eDAVE - extension of GDC Data Analysis, Visualization, and Exploration Tools"

### Slide 1
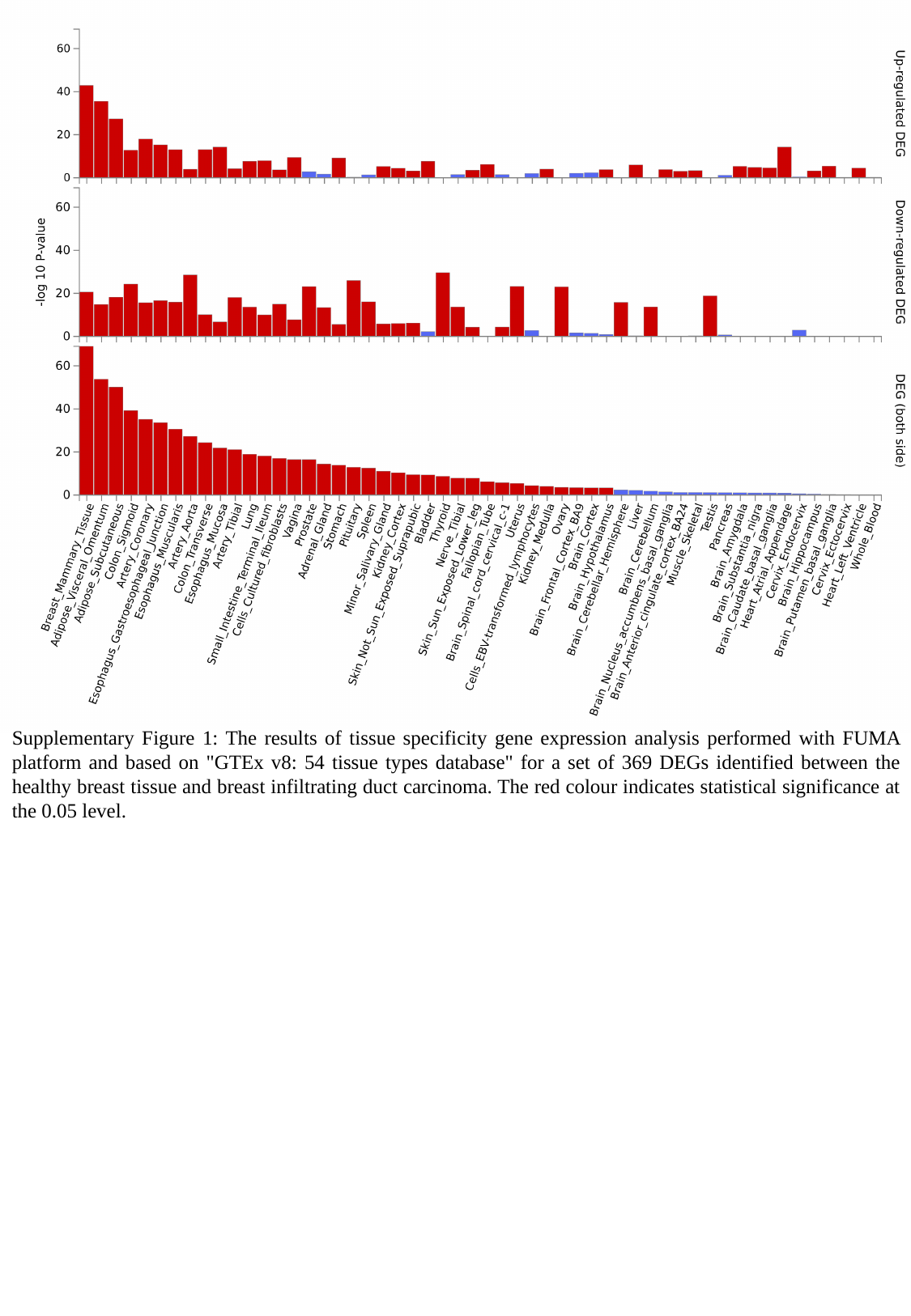

Supplementary Figure 1: The results of tissue specificity gene expression analysis performed with FUMA platform and based on "GTEx v8: 54 tissue types database" for a set of 369 DEGs identified between the healthy breast tissue and breast infiltrating duct carcinoma. The red colour indicates statistical significance at the 0.05 level.
